# ChiraKit, an online tool for the analysis of circular dichroism spectroscopy data

**DOI:** 10.1101/2025.01.29.635490

**Authors:** Osvaldo Burastero, Nykola C. Jones, Lucas A. Defelipe, Uroš Zavrtanik, San Hadži, Søren Vrønning Hoffmann, Maria M. Garcia-Alai

## Abstract

Circular dichroism (CD) spectroscopy is an established biophysical technique to study chiral molecules. CD allows investigating conformational changes under varying experimental conditions and has been used to understand secondary structure, folding and binding of proteins and nucleic acids. Here, we present ChiraKit, a user-friendly, online, open-source tool to process raw CD data and perform advanced analysis. ChiraKit features include the calculation of protein secondary structure with the SELCON3 and SESCA algorithms, estimation of peptide helicity using the helix-ensemble model, the fitting of thermal/chemical unfolding or user-defined models, and the decomposition of spectra through singular value decomposition (SVD) or principal component analysis (PCA). ChiraKit can be accessed at https://spc.embl-hamburg.de/.

## 1. INTRODUCTION

Circular dichroism (CD) spectroscopy is a biophysical technique to study chiral molecules. It measures the difference in absorbance between left- and right-handed circular polarised light as it passes through a sample (1). The CD spectrum contains broad structural information. In the case of proteins, different amounts of secondary structure elements, such as alpha-helices, beta-sheets or turns, produce different CD spectra (2). For nucleic acids, the signal is mostly determined by the geometry of the stacked bases (3–5). Benchtop instruments typically measure in the range of 190 to 340 nm, while synchrotron radiation CD (SRCD) beamlines extend the lower limit to 170 nm, for biomolecules in solution (2).

CD measurements are easy, quick, low cost, non-destructive in some cases, and consume small amounts of sample. CD is useful for assessing conformational changes under varying experimental conditions. It has been used to understand protein folding through thermal or chemical denaturation, to analyse DNA-ligand binding, to validate the quality of mutant proteins, or to study reaction kinetics, among others (6–10). The analysis of CD data ranges from simple workflows, encompassing the averaging, subtraction and normalisation of spectra, to the fitting of secondary structure models (for proteins) or models where the CD signal is expressed as a function of a certain experimental parameter (e.g., temperature).

Numerous computational tools have been developed to extract insights from CD data. CDToolX (Windows), enables standard processing and singular value decomposition (11). DichroWeb and BeStSel (websites), calculate protein secondary structure content (12, 13). DichroWeb provides five algorithms (SELCON3, CONTINLL, CDSSTR, VARSLEC and K2D) (14–18). BeStSel was optimised for beta-sheet-rich and has a module for fold recognition (19). SESCA (Python package), predicts CD spectra from structures and estimates secondary structure content (20, 21). CDPal (Windows), can be used to monitor protein thermal and chemical stability (22). CalFitter (website), offers complex models for thermal unfolding studies (23, 24). Lastly, there are two CD databases: the Protein Circular Dichroism Data Bank (PCDDB), and the Nucleic Acid Circular Dichroism Database (NACDB) (25, 26).

In this work, we present ChiraKit, the latest addition to our eSPC data analysis platform for molecular biophysics (27–29). ChiraKit is open-source, online, user-friendly, interactive, free, and provides functions for pre-processing and advanced analysis, including the deconvolution of spectra, thermal and chemical unfolding models, user-defined models, and protein secondary structure calculations. ChiraKit can be accessed at https://spc.embl-hamburg.de/.

## 2. SOFTWARE DESCRIPTION

The usage workflow can be divided into four steps (Figure 1): The data importing step, then the pre-processing, finally the analysis and export. To start with, the raw CD data is processed and the final spectra are obtained. Secondly, the CD data is analysed as a function of a certain experimental parameter, or the protein secondary structure is calculated. Models for thermal and chemical unfolding are included. A module for the comparison of spectra is also available. Lastly, the final spectra, the fitted parameters, the fitted curves and a logbook containing all the performed steps can be exported.

**Figure 1.**
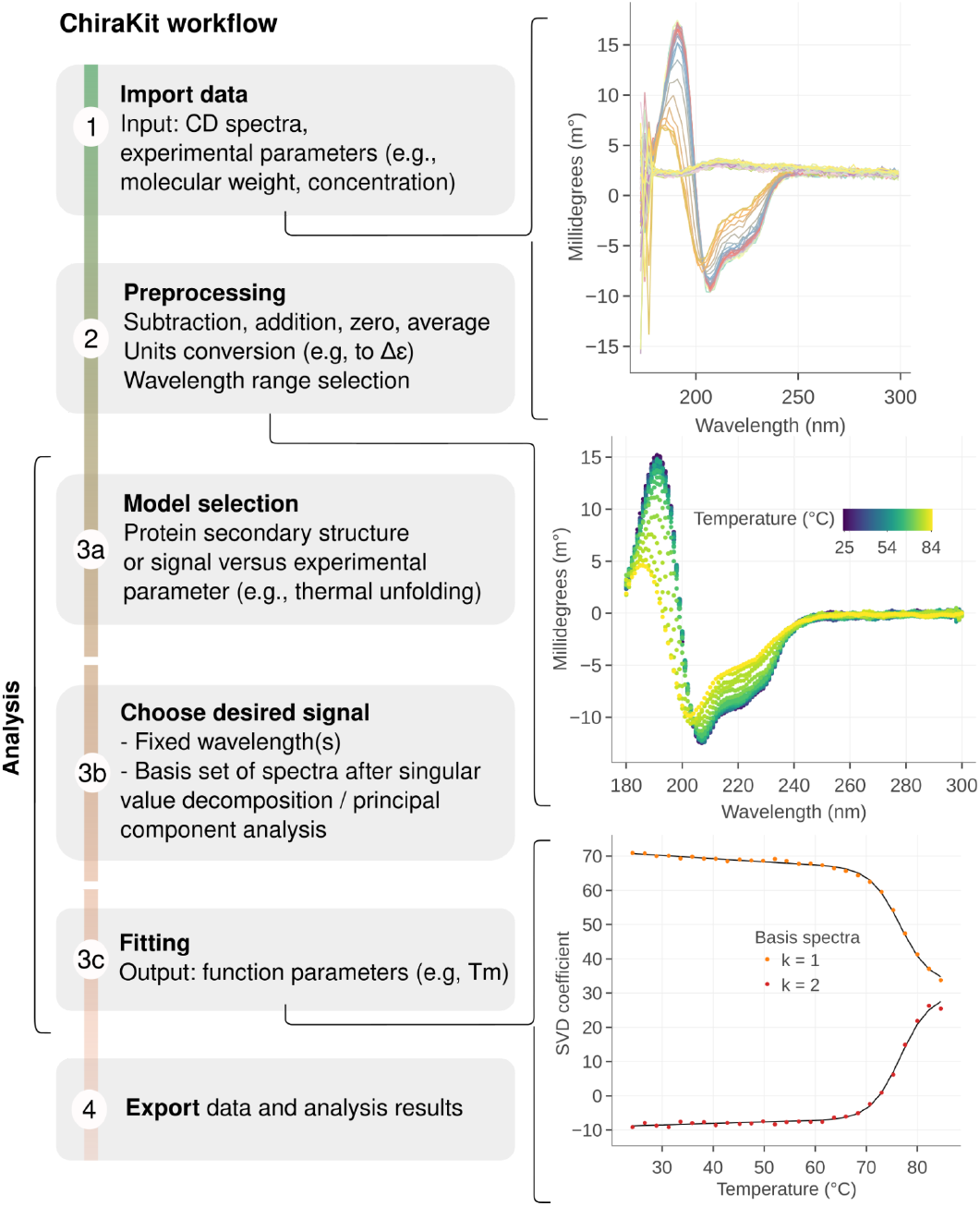
Usage workflow of ChiraKit. Firstly, the data file is imported and the experimental parameters, such as path length, are inputted (1). Subsequently, the CD units are selected (e.g., molar extinction) and the spectra are pre-processed (2). Thirdly, the CD spectra are analysed. For proteins, the secondary structure can be estimated, or, in the case of thermal/chemical unfolding experiments, two or three-state models can be applied. For other types of biomolecules (or experiments), the CD signal can be investigated by employing user-defined models. Moreover, the analysis can focus on single wavelengths or use a set of basis spectra derived from singular value decomposition / principal component analysis (3b). After selecting a model, the fitting is done and the relevant parameters such as the temperature of melting (*T*_m_) are obtained (3c). Lastly, the finalised spectra, fitted parameters and fitted curves can be exported, in text format and/or as figures (if applicable) (4).

### 2.1. Input file

ChiraKit accepts several file types, including those generated by ‘Applied Photophysics’ and ‘JASCO’ instruments, generic (.gen) files for PCDDB/NACDDB databases, PCDDB (.pcd) files, SRCD (.dx or .dat) files, and CSV files with column-formatted data. High-tension voltage data, associated with the CD data collected, if available, is also parsed and displayed.

### 2.2. Processing

CD spectra can be added, subtracted, baseline corrected (zeroed), smoothed and averaged. Six working units are available: millidegrees, differential absorbance, molar extinction, molar ellipticity, mean unit molar extinction (Δε), and mean unit molar ellipticity ([Θ]). Molar extinction/ellipticity units normalise the CD data to the path length, concentration, and molecular weight. [Θ] and Δε are useful for proteins because they take into account the number of chromophores (e.g., peptide bonds) (30). The wavelength range can be manually adjusted or automatically by using a high-tension voltage threshold.

### 2.3. Analysis options

To calculate a protein’s secondary structure content, we provide the SESCA bayesian method (21) and a Python-based implementation of the SELCON3 algorithm (31). These methods are well-suited for proteins, but not for completely unstructured proteins or peptides. The default reference sets for SELCON are based on the SP175 (32) and SMP180 (33) sets, but custom reference sets can be imported. For peptides adopting only helix or coil conformations, the helicity estimation using the ensemble model by Zavrtanik *et al*., 2024, is available (34).

Multiple spectra can be reduced to a set of basis spectra. The basis spectra can be linearly combined to reconstruct all the original spectra. Hence, we take advantage of the information provided by the full spectrum rather than a single wavelength. Two methods are provided: Singular value decomposition (SVD) and Principal component analysis (PCA) (without Z-score normalisation). PCA is equivalent to SVD after centering the data (35). The number of relevant basis spectra is determined by the user, and each basis spectrum will be associated with as many coefficients as the original spectra (e.g., one coefficient per temperature). To help interpreting the results, the spectra can be inverted and/or rotated. The inversion operation changes the sign, while the rotation operation consists of creating a new basis set with one basis spectrum that resembles the first measured spectrum (e.g., lowest temperature) (36).

For protein unfolding analysis, the user can fit two- and three-state reversible unfolding models for chemical and thermal unfolding (37). In the former case, the models are based on the empirical Linear Extrapolation Model (LEM) and depend on the parameters *D50* and *m*, where *D50* is the denaturant concentration at which the free energy equals zero, and *m* is a parameter that correlates with the solvent accessible area (38, 39). In the latter case, the two important thermodynamic parameters are the enthalpy of unfolding (Δ*H*) and the temperature where the free energy equals zero (temperature of melting *T*_m_ or *T*_1/2_).

ChiraKit can fit many curves simultaneously with shared thermodynamic parameters. For instance, the CD signal monitored at wavelengths 198 and 222 nm must have the same *T*_m_ but may have different baselines. The global fitting can be done at the protein concentration level, improving the estimation of dimer, trimer and tetramer unfolding models. Furthermore, two and three-state unfolding models that lead to an irreversibly denatured state (Lumry-Eyring) are available for melting curves of monomers. (40). The two additional important thermodynamic parameters are the energy of activation (*E*_a_) and the temperature where the rate constant equals one (*T*_f_).

The CD signal can be also studied as a function of any desired experimental parameter. The user needs to associate each spectra with a certain value of the experimental parameter and input a fitting function as a string. This approach could be relevant for other types of CD datasets such as 2,2,2-trifluoroethanol (TFE) titrations or DNA-ligand binding (8, 41, 42).

Lastly, ChiraKit has a module to compare samples (e.g., wild-type versus mutant) or the same sample at different conditions. Given two or more groups of spectra, the averages and associated standard deviations are computed, together with the ‘difference’ spectrum (of the averages). To assess the importance of the differences, the intra and inter groups euclidean distances are provided. To compare the shapes of spectra and discard the influence of differences in absorbance intensity, spectra can be normalised to have unit length of one (L2 normalisation) (43).

### 3.1. CASE STUDIES

#### Secondary structure and thermal unfolding of hen egg-white lysozyme

Lysozyme, an enzyme known for hydrolyzing bacterial cell wall peptidoglycan, is a model protein for studies on stability and unfolding. Hen egg-white lysozyme solubilized in water was measured using SRCD. Six scans were performed for both lysozyme in water and water alone. Following data acquisition, the averaged water spectrum was subtracted from the averaged sample spectrum. The CD signal was zeroed using the mean signal in the 296–300 nm range, and the lower wavelength limit was set to 176 nm, based on a voltage threshold indicating low levels of light. The final CD spectrum was normalised to Δε units. The processing was done with the ChiraKit Processing submodule (under 1. Import data).

The secondary structure content was assessed using the SELCON3 method with the ‘AU-SP175’ reference set (Hoffmann, S.V., Jones, N.C., Rodger, A. QRB Discovery (2025) Accepted) (ChiraKit module ‘2c. Protein Secondary Structure’). The calculated values confirm lysozyme’s predominantly alpha-helical structure (40 versus 41.4 %) (Table S1). This outcome is expected, given that lysozyme’s spectrum is included in the Selcon3 reference set. The beta-sheet content differs by a factor of two, highlighting the deviations of the method (12 versus 6.3 %). Furthermore, we applied the SESCA method with the ‘DSSP-TSC1’ basis set, yielding values consistent with the crystallographic structure (Table S2), as anticipated because lysozyme is part of the training set. An advantage of SESCA is that it provides error estimates for each secondary structure element.

To investigate the thermal unfolding of lysozyme, a thermal ramp from 24 to 85ºC at an average heating rate of 1 ºC/minute was performed. Single scans were taken at each temperature step and spectra of water measured under the same conditions as the sample was used as a baseline reference at each temperature. The CD spectra generally reduce in magnitude with increasing temperature, with a change in shape, whereby the peak maxima shifts to lower wavelengths at higher temperature (Figure 2A). The presence of an isodichroic point between 203 and 204 nm suggests that the unfolding process involves two states. If that is the case, fitting the CD signal at any particular wavelength should yield similar thermodynamic parameters.

**Figure 2.**
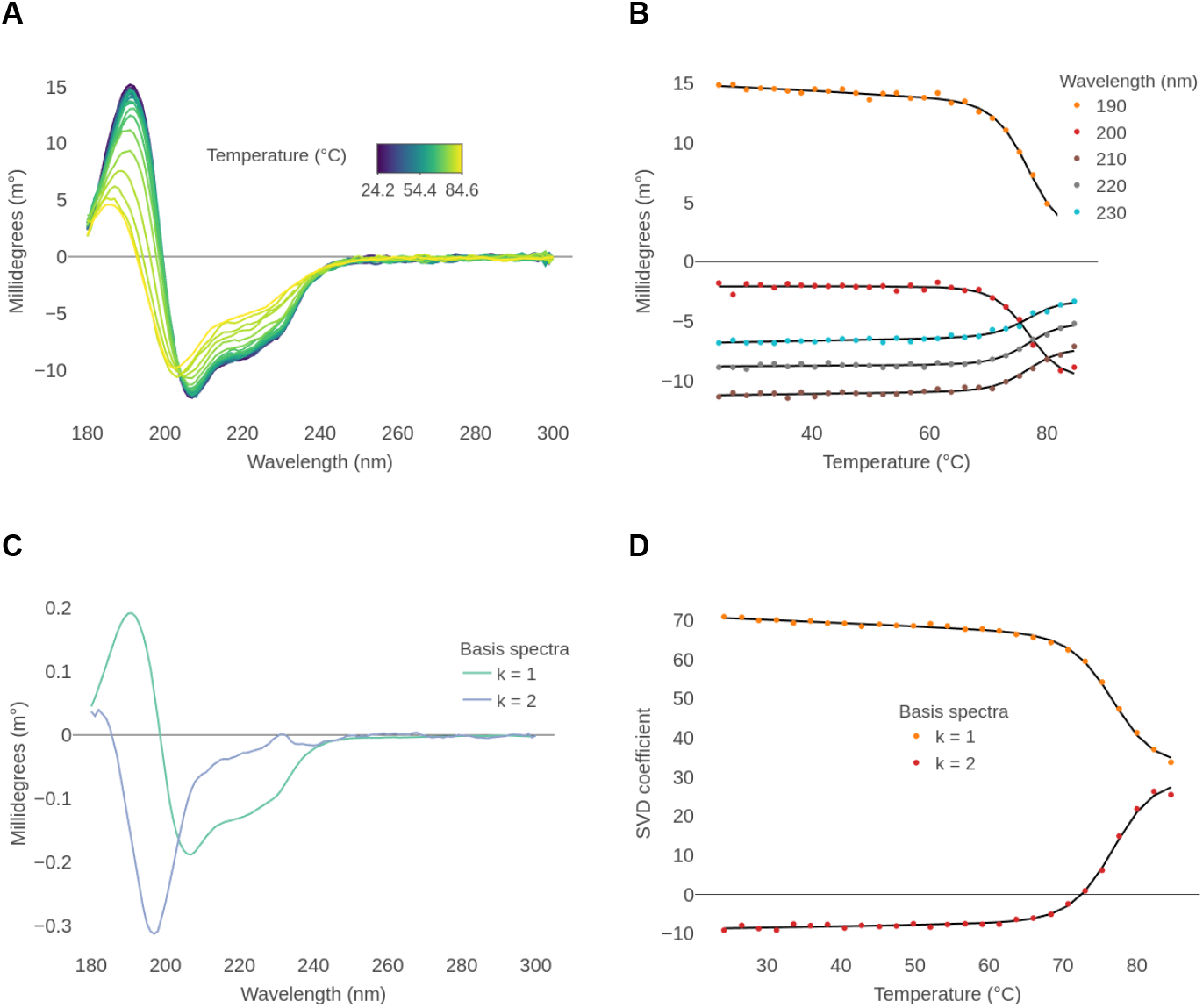
Two-state thermal unfolding analysis of lysozyme. A) Finalised spectra after baseline subtraction and wavelength range correction. B) Points: CD signal versus temperature, at five selected wavelengths. Lines: global fitted values based on a two-state reversible unfolding model. Baseline and slope (folded state only) values vary freely for each curve, while the parameters *T*_m_ and Δ*H* are shared. Δ*Cp* was set to 2.3 kcal/K/mol. C) SVD derived basis spectra required to reconstruct the original spectra. The cumulative explained variances are 97 and 99.93 %. A rotation was performed such that the first basis spectrum resembles the spectrum measured at the lowest temperature. D) Points: SVD coefficients versus temperature, for the two basis spectra. Lines: global fitted values based on a two-state reversible unfolding model. Baseline and slope (folded state only) values vary freely for each curve, while the parameters *T*_m_ and Δ*H* are shared. Δ*Cp* was set to 2.3 kcal/K/mol.

We applied a global fitting at different wavelengths (190 to 230 nm, every 10 nm) and obtained a *T*_m_ of 76.6 ± 0.2 ºC and Δ*H* of 86.8 kcal/mol ± 3.5 kcal/mol (Figure 2B), in agreement with previous estimations (44, 45) (ChiraKit module 2a. Thermal unfolding). We recommend considering the relative errors of the fitted parameters as a criterion for discarding a model. These optimistic errors, as in almost all other non-linear fitting software, are calculated from the covariance matrix of the fitted parameters and assume a linear model.

An improved approach to fully use the entire wavelength range information is spectral decomposition. Reconstructing the data post-SVD decomposition requires two basis spectra (Figure 2C). Global fitting of the SVD-associated coefficients yields highly consistent estimates: a *T*_m_ of 76.6 ± 0.1 °C and a Δ*H* of 88.1 ± 3.5 kcal/mol (Figure 2D).

#### Secondary structure and chemical unfolding of the Clathrin Heavy Chain N-terminal domain

The clathrin heavy chain N-terminal domain (CHC-NTD) serves as the primary contact point between clathrin and adaptor proteins, linking the clathrin cage to the membrane through the binding of short linear motifs in the adaptor proteins (46). It consists of seven beta-transducin (WD) repeats that form a beta-propeller, resulting in a predominantly beta-sheet structure with two short alpha helices (see Figure S1). The CD spectrum of the CHC-NTD solubilized in sodium phosphate buffer with sodium fluoride salt was measured using a CD benchtop instrument. The lowest, noiseless, wavelength was 188 nm, and the highest wavelength was

300 nm. After averaging the scans and subtracting the baseline, the CD spectrum was converted to Δε (mean unit molar extinction) and the secondary structure was calculated with the Selcon method. The expected values are in agreement with the mixed beta/alpha structure, although the absolute percentages of beta and alpha deviate almost 1.6-fold from the values obtained via DSSP analysis of the crystal structure (Table 1).

**Table 1.** Comparison of Clathrin Heavy Chain N-terminal domain secondary structure composition as determined by the Selcon3 method versus structural data.

| Method | Alpha | Beta | Turns | Other |
| --- | --- | --- | --- | --- |
| SELCON | 14 | 33 | 15 | 36 |
| DSSP (PDBid 5m5r chain A) | 8.3 | 51.4 | 15.6 | 24.7 |

Then, CD spectra of the CHC-NTD solubilized in Tris buffer with sodium chloride salt were acquired. The urea concentration ranged from 0.5 to 6.5 M, with steps of 0.5 M. Scans taken at the same urea concentration were averaged. The spectra were recorded up to 260 nm and the buffer spectrum was used as a baseline reference. To remove noisy data, the lower wavelength was set to 212 nm. Concentrations of urea ranging from 0.5 to 3.5 were measured in triplicate, while concentrations from 4 to 6.5 M were measured in duplicate, yielding 33 finalised spectra (Figure 3A).

**Figure 3.**
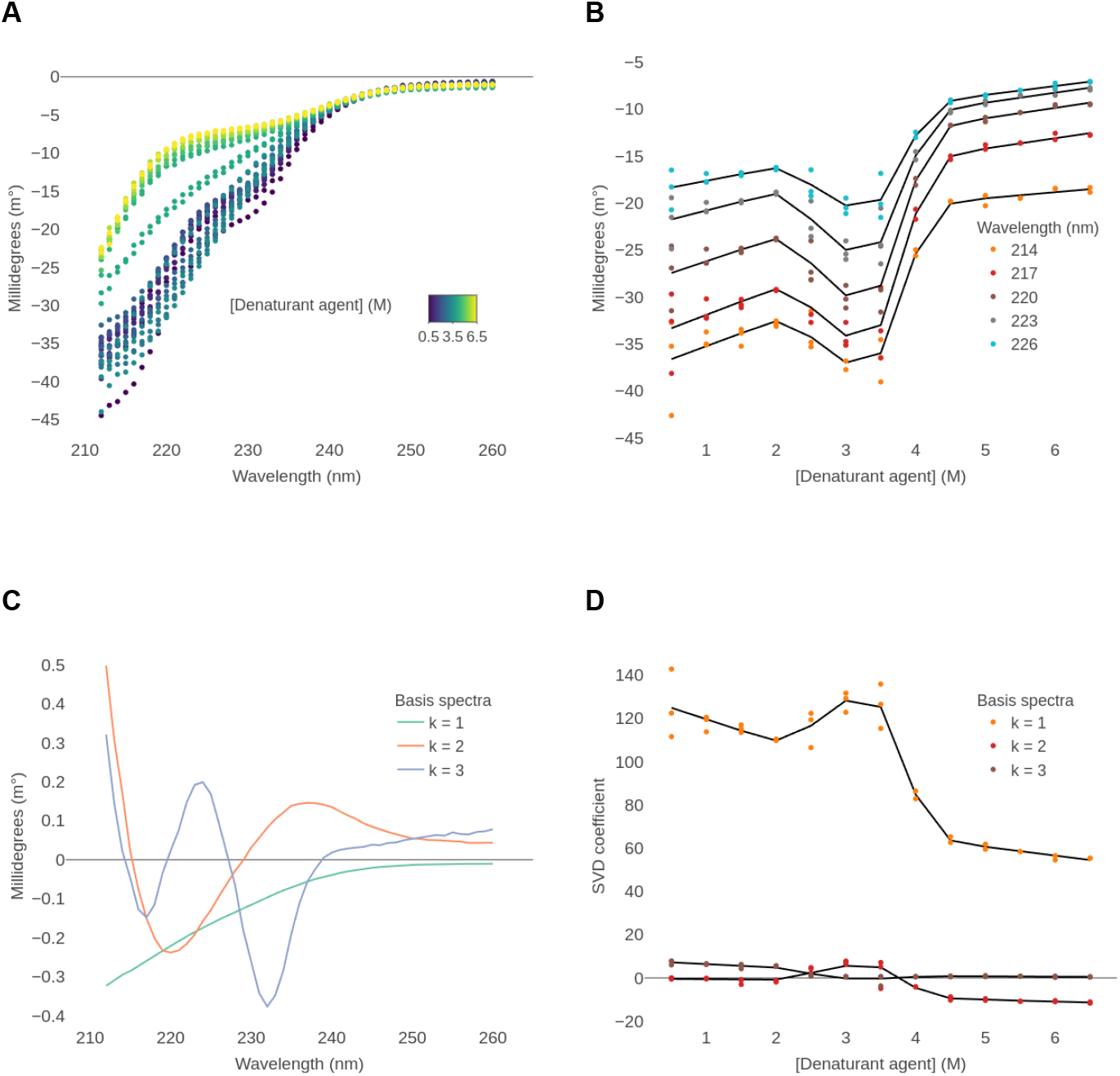
Three-state chemical unfolding analysis of the clathrin heavy chain N-terminal domain. A) Finalised spectra after baseline subtraction and wavelength range correction. B) Points: CD signal versus urea concentration, at five selected wavelengths. Lines: global fitted values based on a three-state reversible unfolding model. Baseline and slope values vary freely for each curve, while the parameters *D50*_1_, *D50*_2_, *m*_1_ and *m*_2_ are shared. C) SVD derived basis spectra required to reconstruct the original spectra. The cumulative explained variances are 99.43, 99.84 and 99.97 %. A linear rotation was performed such that the first basis spectrum resembles the spectrum measured at the lowest urea concentration. D) Points: SVD coefficients versus v, for the three basis spectra. Lines: global fitted values based on a three-state reversible unfolding model. Baseline and slope values vary freely for each curve, while the parameters *D50*_1_, *D50*_2_, *m*_1_ and *m*_2_ are shared.

A visual inspection of the spectra reveals a complex unfolding pattern, with the CD signal initially reducing in magnitude with an increasing concentration of urea up to 2 M. Then the CD signal increases again to a maximum around 3-3.5 M, before decreasing again to higher concentrations. This effect can be appreciated in Figure 3B, where single wavelength data is shown. Considering the two transitions, one between 2 and 3 M urea, and the other between 3.5 and 5 M urea, we applied a three-state reversible unfolding model (N ⇋ I ⇋ U) (ChiraKit module 2b. Chemical unfolding).

We analysed both the SVD coefficients and single wavelength data (Figure 3). The inclusion of the pre- and post-transition slopes resulted in a better fitting. The estimated values of *D50*_1_ and *D50*_2_ were 2.5 and 3.9 M (Table S3 and S4). These parameters are the urea concentrations at which Δ*G*_1_ (N ⇋ I) and Δ*G*_2_ (I ⇋ U) equal zero, respectively. The *m*-value of the first transition could not be reliably estimated, while the second *m*-value was ≈ 4 kcal/mol/M. The *m*-values are linearly related to the difference in solvent accessible surface area between the native and the unfolded states (39, 47, 48). Indeed, the predicted m-value according to the number of residues for the global unfolding process would be ≈ 4.5 kcal/mol/M. The second *m*-value is similar to the global *m*-value, suggesting that the most significant structural changes occur during this step.

#### Analysis of transient helicity in intrinsically disordered proteins

Intrinsically disordered proteins (IDPs) lack a fixed three-dimensional structure under physiological conditions. Nevertheless, they can transiently sample elements of secondary structure or acquire structure upon binding their partner proteins (49). Common interaction motifs are the helical binding motifs, short IDP segments that fold into an α-helix upon binding (50). When these regions are unbound, the helical structure is partially formed, and overlaps with the target-binding motif (51). CD spectra enable quantification of helix content in IDP peptides undergoing folding-upon-binding. Since peptides exist as conformational ensembles, helicity cannot be estimated using algorithms such as SESCA. A traditional method is based on a simple algebraic relation that normalizes the measured signal at 222 nm to the ellipticity of maximal helix (52). The drawback is that, since ellipticity depends on the length of the helix segment (53), it performs well only for peptides whose ensembles consist of conformers with long helical segments, approaching maximal helix length. In contrast, IDP ensembles typically consist of conformers with short or intermediate helices due to their lower helix propensity, leading to underestimation of helix content. The developed helix ensemble model (34), integrated in ChiraKit, estimates the average peptide helicity by summing together the individual length-corrected contributions of all conformers in the peptide ensemble.

The CD spectra of a set of IDP peptides (Table S5) that fold-upon**-**binding was measured using a benchtop instrument in phosphate buffer (Figure 4A). Prior to analysis, spectra were buffer subtracted, and CD intensity was calibrated using camphor sulfonic acid. The intensity at 222 nm was then used to estimate fractional helicity (*f*_H_) content, with obtained *f*_H_ values ranging between 0.1 to 0.3 (Table S6). For comparison, we show that helicity estimated using the conventional approach strongly underestimates peptide helicity for IDPs (Table S6, CcdA, error more than 200%) but performs rather well for peptides with high helix tendency (Lifson-Roig *w* > 1.2 (54)), such as alanine peptides (AK32, error around 5%). Note that the helix ensemble model assumes only an equilibrium between coil and helix conformations; therefore, it is not suitable for analysis of peptides exhibiting β-sheet propensity.

**Figure 4.**
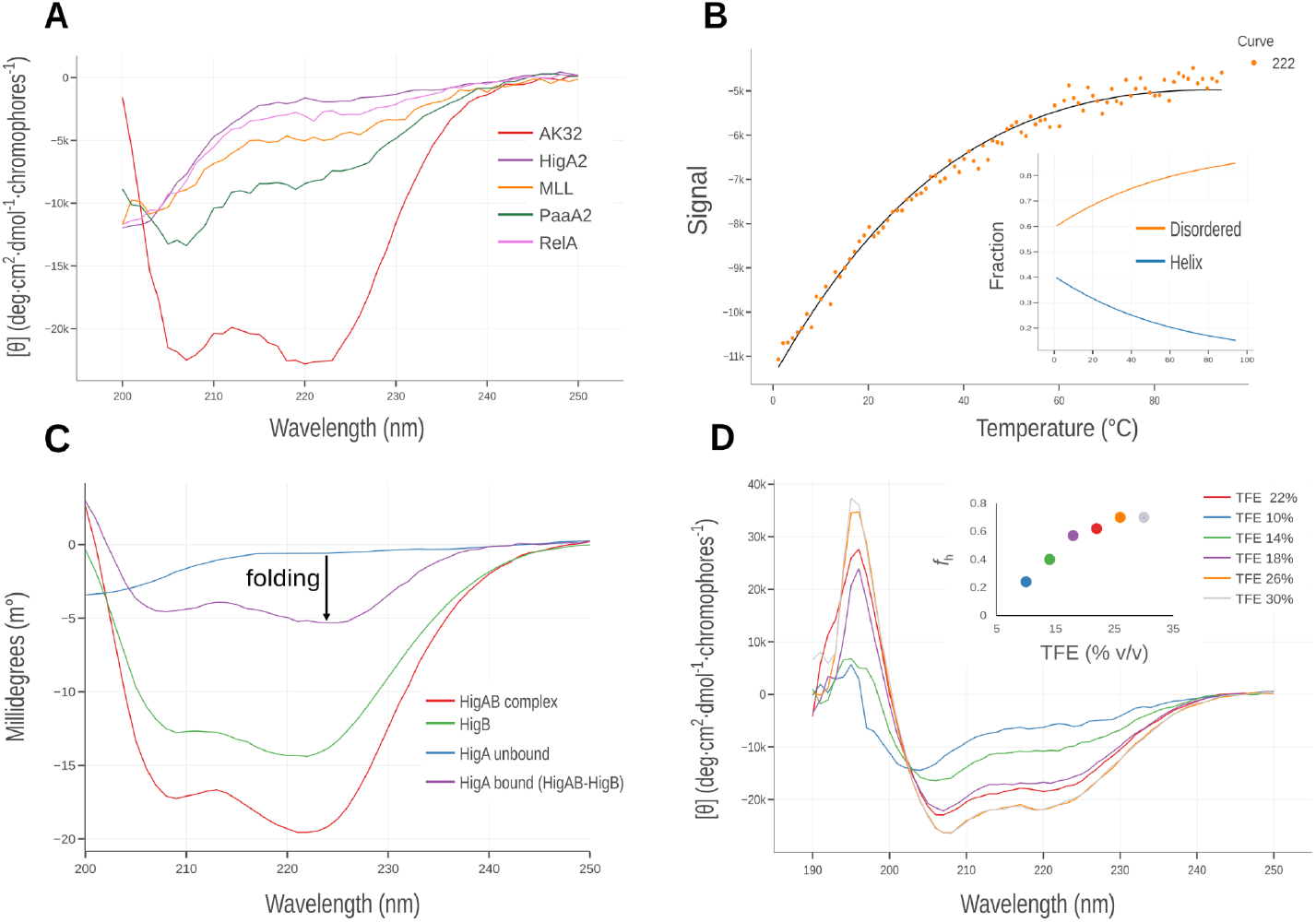
A) Far-UV CD spectra of intrinsically disordered peptides and alanine-rich peptide (red). Mean residue ellipticity at 222 nm and the corresponding helix fraction (*f*_H_) estimated helicity are reported in Table S6. B) Thermal denaturation of the PaaA2 peptide monitored by CD intensity at 222 nm. Orange points represent experimental data, and the solid black line shows the model fit using the ensemble model. The inset displays the ensemble model-derived overall helicity as a function of temperature. C) Folding-upon-binding analysis of the intrinsically disordered antitoxin HigA upon binding to its globular target toxin HigB. The difference spectrum of the target-bound antitoxin HigA (purple) was obtained by subtracting the spectrum of the free toxin (HigB, green) from that of the HigBA complex (red). D) TFE induction experiment of the disordered HigA peptide. The inset shows helicity as a function of the volume percentage of TFE.

Thermal denaturation can provide insights into forces stabilizing helix structure and help with the energetic dissection of folding and binding contributions of IDP-target interactions. For the PaaA2 peptide, with considerable helicity (*f*_H_ > 0.2), we measured intensity at 222 nm as a function of temperature (Figure 4B). Differently from proteins, where thermal denaturation induces highly cooperative transitions, in the case of peptides, loss of helix content occurs with low cooperativity. We used the helix ensemble model to analyze PaaA2 data available under ‘2.a. Thermal unfolding’ module and obtained thermodynamic parameters of transition Δ*G* = -0.07 and Δ*H* = -0.3 kcal mol^-1^ pep. bond^-1^ (Figure 4B). Assuming that 15 residues of PaaA2 fold into α-helix upon binding its target (55), the associated Δ*H*_folding_ for PaaA2 can be estimated to be around 5 kcal mol^-1^.

CD spectroscopy is an excellent tool to study IDP folding-upon-binding processes. For example, intrinsically disordered HigA folds into an α-helix upon binding the globular target HigB (56). This process can be studied by measuring CD spectra of unbound HigA2 and HigB2 and a 1:1 HigA2-HigB2 complex (Figure 4C). The ChiraKit Processing submodule (under 1. Import data) can then be used to obtain a difference spectrum (i.e. complex-HigB2), corresponding to folded HigA2, assuming no structural change in HigB2. Comparison with the unbound state (Table S6) then shows that helix content in HigA2 increased from *f*_H_ = 0.12 to about *f*_H_ = 0.7 upon binding HigB2. Finally, in some cases where no binding partner is available, the potential of IDPs to adopt a helix conformation can be assessed using helix-inducing cosolvents. For example, titration with TFE of HigA2 peptide shows considerable gain in helicity which stabilizes around 25% TFE, where it reaches *f*_H_ = 0.7, as calculated using ensemble model (Figure 4D).

#### Re-analysis of published data

To further validate ChiraKit, we reanalyzed three published datasets with known parameters. These include one protein-DNA binding affinity measurement (ChiraKit module 2e. Custom analysis), two-state unfolding of a protein dimer (module 2c. Chemical unfolding) and three-state unfolding of a protein dimer (module 2c. Chemical unfolding). The three cases are presented in the Supporting Information (Figure S2 to S4).

## 4. CONCLUSIONS

In this work, we presented and evaluated ChiraKit, an open-source, free, multi-purpose tool for processing and analysing CD data. ChiraKit supports multiple input file formats, multiple CD units and pre-processing functions. It does not require installation, nor registration and it provides a user-friendly, interactive interface. ChiraKit provides modules for the comparison of spectra, estimation of peptide helicity, estimation of protein secondary structure, and model-based analysis of thermal and chemical unfolding.

Regarding the case studies, the thermal unfolding of hen egg-white lysozyme could be explained with a two-state reversible unfolding model. The chemical unfolding of the Clathrin Heavy Chain N-terminal (CHC) domain follows a three-state process, with major structural changes occuring between the intermediate and final states. The secondary structure estimation of CHC shows an overall agreement with the values extracted from the PDB, with deviations of up to 2-fold.

IDPs may adopt secondary structures, such as α-helices, upon binding to partner proteins. The ChiraKit helix-coil ensemble model provides accurate helicity estimates by considering the contributions of all conformers in peptide ensemble, thus enabling estimation of helical content of unbound IDP and the analysis of the folding-upon-binding process.

In the presented cases, the whole pipeline, from the pre-preprocessing of the CD raw data to the fitting and figure generation, was done in ChiraKit. We expect that CD users find this tool useful and look forward to new developments.

## 5. METHODS

### 5.1. Lysozyme sample origin

Hen egg-white lysozyme was purchased from Sigma Aldrich (CAS-Nummer:12650-88-3). The sample was prepared by dissolving the powder in MilliQ water (no buffer or salts). The concentration of the sample, 1.1 mg/ml, was determined from the measured absorbance at 205 nm using the sequence from the UniProt ID P00698 (UniProt Consortium, 2023). The sample was prepared from the powder and kept in the fridge until the measurements (same day).

### 5.2. Synchrotron radiation circular dichroism (SRCD) measurements of lysozyme

The CD spectra were recorded on the AU-CD beam line of the ASTRID2 synchrotron radiation light source at the Department of Physics and Astronomy, Aarhus University. Correct operation of the CD spectrometer was confirmed through measurement of a known concentration of camphor sulphonic acid (CSA, CAS number: 3144-16-9). Measurements were carried out with the sample in a nominal 0.1 mm path length quartz suprasil cell (Starna type 31B), with the path length determined to be 0.1106 mm. Measurements were carried out using the fast scanning mode, whereby the monochromator is in constant motion through the wavelength range, allowing data to be collected for a single scan in just under 1 minute. One scan was taken at each set temperature step of 2.5°C with an equilibration time of 1 minute after the temperature was reached, before starting each measurement.

### 5.3. Expression and purification of the *Homo sapiens* clathrin heavy chain N-terminal domain (CHC-NTD)

The *Homo sapiens* clathrin heavy chain N-terminal domain (Chc-NTD, 1-364) was cloned in a pETM-30 vector which contains a His-GST tag and a TEV cleavage site. Chemo-competent *E. coli* BL21 (DE3) cells were transformed with 100 ng of plasmidic DNA and grown overnight at 37 °C with 30 µg/ml of kanamycin. For protein expression, a 1:100 dilution of the preculture was done in 2xTY (16 g tryptone, 10 g yeast extract, 5 g NaCl per litre). Cells were grown at 37°C until reaching OD 0.8-1. Then, the temperature was decreased to 18°C, and induced by adding IPTG to a final concentration of 0.3 mM. Induction was performed overnight. Cells were harvested at 12,000 x g for 15 minutes at 18°C and stored at -20°C until purification. Cells were resuspended in 5 to 10 mL of lysis buffer (20 mM sodium phosphate pH 7.5, 200 mM NaCl, 0.05% (w/v) Tween-20, 2 mM MgCl2, 0.5 mM TCEP, +400 U DNAse I, 12.5 mM imidazole and a tablet of Complete EDTA-free protease inhibitor cocktail, Roche, per 100 mL of buffer) per gram of cells. Lysis of the cells was done with an Emulsiflex C3 (Avestin, Ottawa, Canada) cell disruptor at 15 kPsi three times. A centrifugation at 42,000 x g for 1 hour at 4°C was performed to clear the lysate.

The lysate was filtered with a 0.45 µm filter and loaded onto a Ni-NTA (Carl Roth, Germany) gravity column equilibrated with buffer A (20mM sodium phosphate pH 7.5, 500 mM NaCl and 12.5 mM imidazole). The column was washed with 10 column volumes (CV) of buffer A and eluted with buffer B (20mM sodium phosphate pH 7.5, 500 mM NaCl and 250 mM imidazole), collecting 1 mL fractions. The purity of the protein samples was assessed using SDS-PAGE. Subsequently, selected fractions were pooled and treated with TEV protease at a ratio of 1 mg of TEV per 25 mg of protein. This mixture was then diluted in a 1:1 ratio with a dialysis buffer composed of 20 mM TRIS (pH 7.5), 200 mM NaCl, and 0.5 mM TCEP. Finally, the solution was subjected to overnight dialysis at 4°C with the same buffer.

A reverse Ni-NTA was performed to remove HisTag-TEV and the cleaved HisGST from the solution. A prepacked Ni-NTA column was equilibrated with a dialysis buffer and the flowthrough was collected and concentrated to 2 mL using a 10 kDa MWCO concentrator (Amicon Ultra 15, Millipore). The sample was loaded onto a Superdex 200 HiLoad 16/600 size-exclusion chromatography (SEC) column equilibrated with SEC buffer (50 mM Tris pH 9, 150 mM NaCl, 0.5 mM TCEP). Fractions were analysed for purity using SDS-PAGE, pooled, concentrated to 20 mg/ml, flash frozen in liquid nitrogen, and stored at -70°C until used.

### 5.4. CD measurements of CHC-NTD (secondary structure)

The CHC-NTD sample was dialyzed overnight at 4°C against 20 mM sodium phosphate buffer (pH 7.5), containing 150 mM sodium fluoride and 0.5 mM TCEP, using a 10 kDa MWCO Slide-A-Lyzer MINI dialysis device (Thermo Fisher Scientific). The final protein concentration was adjusted to 0.133 mg/mL. CD spectra were acquired in the range of 180-300 nm using a ChiraScan spectrophotometer (Applied Photophysics, Leatherhead, UK), equipped with a Quantum Northwest TC 125 temperature controller (Liberty Lake, Washington, USA), set at 20°C. Measurements were performed in a 300 µL, 1 mm pathlength quartz cuvette (Hellma, Müllheim, Germany). Three scans per spectrum were recorded, each with a step size of 1 nm and a dwell time of 1 second.

### 5.5. CD measurements of CHC-NTD (chemical unfolding)

CHC-NTD solutions at a concentration of 0.41 mg/ml were incubated overnight with SEC Buffer or SEC Buffer supplemented with increasing concentrations of Urea, ranging from 0.5 M to 6.5 M in 0.5 M increments. CD spectra were recorded between 205 nm and 260 nm using a ChiraScan spectrophotometer (Applied Photophysics, Leatherhead, UK) equipped with a Quantum Northwest TC 125 temperature controller (Liberty Lake, Washington, USA), maintained at 20°C. Measurements were conducted in a 300 µL quartz cuvette with 1 mm path length (Hellma, Müllheim, Germany). Concentrations from 0.5 to 3.5 M were measured in triplicate, while concentrations from 4 to 6.5 M were measured in duplicate. Three scans per spectrum were recorded, each with a step size of 1 nm and a dwell time of 1 second.

### 5.6. CHC-NTD solvent accessible area surface and m-values

The change in solvent-accessible surface area (Δ*ASA*) of CHC-NTD was calculated using the Equation located in the inset of Figure 1 from ref. (39):

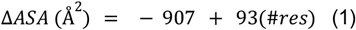

where #*res* is the number of residues (364). The global m-value (urea) was estimated according to the Equation 29.9 of (47):

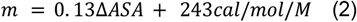

### 5.7. CD measurements of intrinsically disordered peptides

Peptides were purchased from Sigma-Aldrich or China Peptides (AK32 peptide) with a minimum purity of 95%. The sequences are listed in Table S5. Peptides were dissolved and dialyzed against a 20 mM phosphate buffer (pH 7.0) containing 150 mM NaCl, except for the alanine peptide AK32, which was dialyzed in the same phosphate buffer with a higher salt concentration (1 M NaCl). Prior to each experiment, peptide stock solutions were extensively centrifuged, and their concentrations were determined by measuring absorbance at 280 nm using extinction coefficients of 1.450 M−^1^cm−^1^ for tyrosine and 5.690 M−^1^ cm−^1^ for tryptophan (57).

CD spectra were recorded using a JASCO J-1500 spectrometer. All peptides were diluted to a final concentration of 60 µM, except for the AK32 peptide, which was prepared at 100 µM. Spectra were measured in a 1 mm (pathlength) cuvette at 25 °C. The spectrometer was calibrated prior to each measurement using ammonium-d-camphor-10-sulfonate (Sigma-Aldrich). For TFE induction experiments, TFE (trifluoroethanol) from Sigma-Aldrich was used. For thermal melting experiments, ellipticity at 222 nm was monitored at 1°C intervals with an integration time of 8 s per measurement.

## Supplementary Data

Supplementary data are available at X Online.

## Software availability

ChiraKit is available at https://spc.embl-hamburg.de. This website is free and open to all users and there is no login requirement. The source code is freely available at https://github.com/SPC-Facility-EMBL-Hamburg/eSPC_biophysics_platform.

## Acknowledgments

We acknowledge technical support by the Sample Preparation and Characterisation (SPC) facility at EMBL Hamburg, Germany, and the Synchrotron Radiation Circular Dichroism Facility at ASTRID2 (AU-SRCD), Aarhus University, Denmark. We thank the users of the SPC and AU-SRCD for constructive feedback.

## Author contributions

Osvaldo Burastero: Conceptualization, Software (ChiraKit programming), Writing—original draft, Writing—review & editing. Nyk Jones: Data acquisition (CD data of lysozyme), Software (feedback), Writing—review & editing. Lucas A. Defelipe: Data acquisition (CD data of clathrin heavy chain), Writing—review & editing. Uroš Zavrtanik: Data acquisition (CD data of IDPs), Software (feedback), Writing—review & editing. San Hadži: Supervision, Funding acquisition, Software (feedback), Writing—review & editing. Søren Vrønning: Supervision, Funding acquisition, Software (feedback), Writing—review & editing. Maria M. Garcia-Alai: Conceptualization, Supervision, Funding acquisition, Writing—original draft, Writing—review & editing.

## Large language models

ChatGPT (https://chat.openai.com/) was used as an aid to correct written text. ChatGPT and Github Copilot (https://github.com/features/copilot) were used for coding assistance. The authors take full responsibility for the manuscript content and code.

