## Supplementary data for "ChiraKit, an online tool for the analysis of circular dichroism spectroscopy data"

### Supplementary Figures

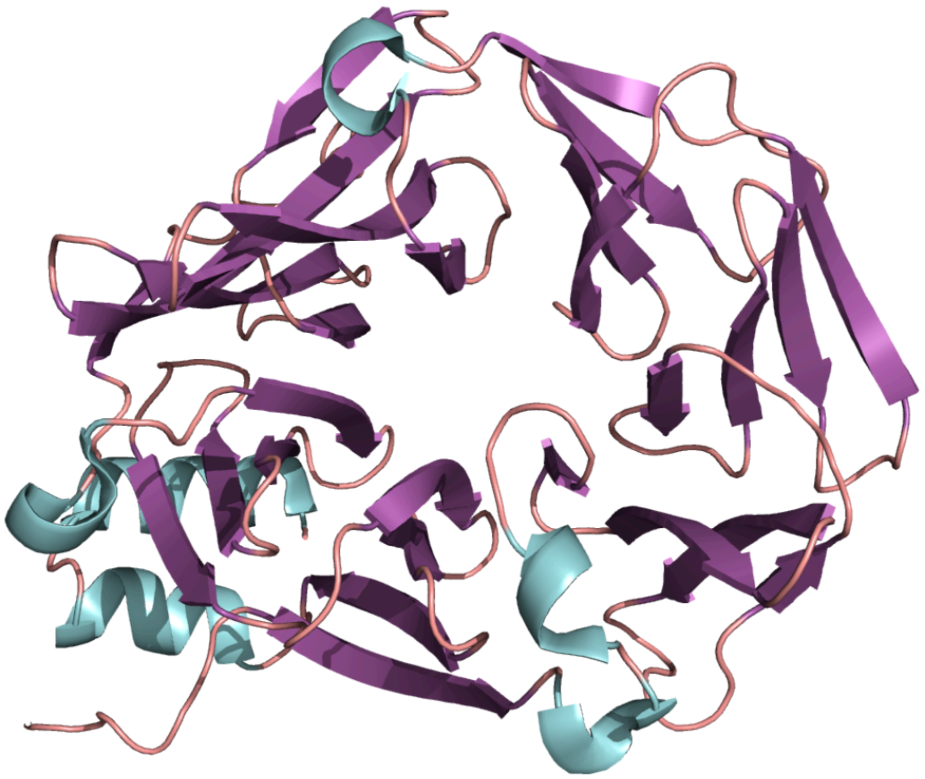

**Figure S1.** X-ray structure of the human Clathrin Heavy Chain N-Terminal Domain (HsCHC-NTD, PDB code: 9C0Y). The structure is shown in cartoon representation, with alpha helices colored in cyan, beta sheets in purple, and loops in salmon.

### Supplementary Tables

**Table S1.** Comparison of lysozyme's secondary structure composition as determined by the SELCON3 method versus structural data.

| Method | Alpha helix | Beta sheet | Turns | Other |
|---|---|---|---|---|
| SELCON3* | 40 | 12 | 15 | 34 |
| DSSP (PDBid 1dpx) | 41.4 | 6.3 | 23.4 | 28.9 |

*The ‘alpha-regular’ and ‘alpha-distorted’ secondary elements were grouped into the ‘alpha’ category. The ‘beta-regular’ and ‘beta-distorted’ secondary elements were grouped into the ‘beta' category.

**Table S2.** Comparison of lysozyme's secondary structure composition as determined by the SESCA bayesian method versus structural data.

| Method | Alpha helix | Beta sheet | Coil |
|---|---|---|---|
| SESCA | 37 ± 6 | 9 ± 9 | 53 ± 7 |
| DSSP-T (PDBid 1dpx) | 41.4 | 6.2 | 52.3 |

**Table S3.** Fitted parameters of the three-state **reversible** model applied to the chemical unfolding of clathrin heavy chain N-terminal domain (CHC-NTD). The first three SVD coefficients were used for the fitting.

| **Parameter** | **Lower fitting bound** | **Fitted value** | **Upper fitting bound** | **Relative error (%)** |
|---|---|---|---|---|
| M1 (kcal/mol/M) | 0.2 | 3.89 | 20 | 148 |
| $D50_1$ (M) | 0.75 | 2.5 | 4.25 | 3.1 |
| M2 (kcal/mol/M) | 0.2 | 4.07 | 20 | 43.2 |
| $D50_2$ (M) | 1 | 3.88 | 6 | 1.8 |

* Rows marked in red contain parameters with relative errors larger than 95 %.

**Table S4.** Fitted parameters of the three-state **reversible** model applied to the chemical unfolding of clathrin heavy chain N-terminal domain (CHC-NTD). The CD signal at wavelengths 214, 217, 220, 223, and 220 nm were used for the fitting.

| **Parameter** | **Lower fitting bound** | **Fitted value** | **Upper fitting bound** | **Relative error (%)** |
|---|---|---|---|---|
| M1 (kcal/mol/M) | 0.2 | 4.72 | 20 | 163 |
| $D50_1$ (M) | 0.75 | 2.49 | 4.25 | 1.7 |
| M2 (kcal/mol/M) | 0.2 | 4.01 | 20 | 27.1 |
| $D50_2$ (M) | 1 | 3.88 | 6 | 1.2 |

* Rows marked in red contain parameters with relative errors larger than 95 %.

**Table S5.** Peptide sequences for the study of intrinsically disordered proteins.

| **Peptide Name** | **Sequence** |
|---|---|
| c-Myb | Ac-EKRIKELELLLMSTENELKGY-NH2 |
| MLL | Ac-PSDIMDFVLKNTPY-NH2 |
| PaaA2 | DPRPAIPHDEVERRMAERFAKMRKERSKQW |
| RelA | DDRHRIEEKRKRTYETFKSIMKKS |
| CcdA | RRLRAERWKAENQEGMAEVARFIEMNGSFADENRDW |
| HigA2 | NRDLFAELSSALVEAKQHSEGW |
| AK32 | Ac-$(AAKAA)_6$GY-NH2 |

**Table S6**. Helix contents of intrinsically disordered peptides with helix binding motifs. From the measured CD signals [θ] we estimated fractional peptide helicity $f_H$ using the ensemble model (1) integrated in ChiraKit (fifth column). This is compared to the traditional approach to estimate $f_H$ (2) (sixth column), which gives large error in case of disordered peptides, but is rather accurate for peptides with high helix propensity (eg. alanine peptide, AK32). LR parameter gives a measure of helix propensity. All data were recorded at 25°C.

| Peptide name | $N_{pep.bonds}$ | [θ] (deg $cm^2$ $dmol^{-1}$ pep.bonds$^{-1}$) | LR propagation parameter - *w* | $f_h$ (ensemble) | $f_h$ = ([θ] - $[θ]_c$) / $Δ[θ]_{h-c}$ | Rel_error (%) |
|---|---|---|---|---|---|---|
| c-Myb | 23 | -4699.1 | 1.06 | 0.21 | 0.13 | 58 |
| MLL | 15 | -5001.6 | 1.20 | 0.23 | 0.16 | 48 |
| PaaA2 | 29 | -7841 | 1.08 | 0.3 | 0.22 | 34 |
| RelA | 23 | -2707.8 | 1.00 | 0.14 | 0.07 | 100 |
| CcdA | 35 | -1568.8 | 0.92 | 0.1 | 0.03 | 210 |
| HigA2 | 21 | -1919.3 | 0.98 | 0.12 | 0.05 | 162 |
| AK32 | 33 | -22574 | 1.27 | 0.7 | 0.66 | 6 |

**Supplementary Results**

**CASE STUDIES WITH PUBLISHED DATA**

**Protein - DNA binding affinity**

The interaction of berenil with calf thymus DNA was reported by Garbett, Nichola C. *et al.*, 2007 (3). We digitised the data from Figure 1B, which represents the CD signal (in millidegrees) at 385 nm versus DNA concentration, and analysed it using the 'Custom analysis' panel (See Supplementary Methods). We applied a binding model that assumes DNA does not contribute to the CD signal, and different signal intensities between the bound and free states of berenil. The fitted $K_D$ was 78 µM, in agreement with the reported one (50 µM) (Figure S1).

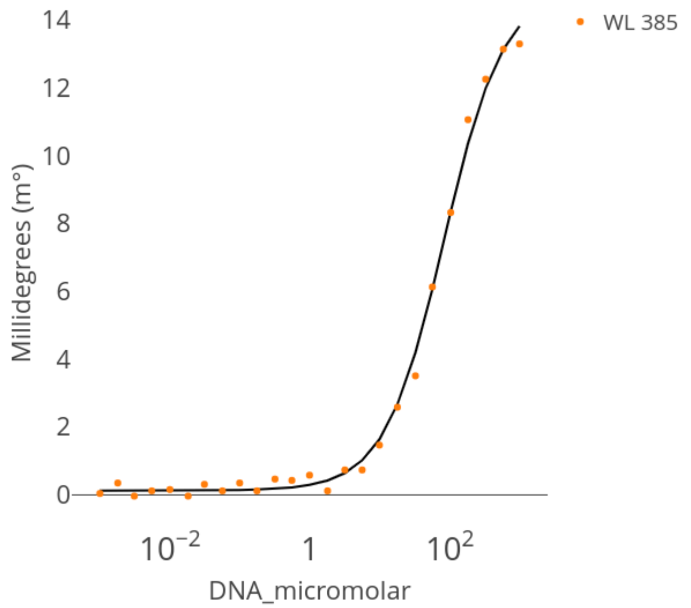


**Figure S2.** CD intensity (at 385 nm) versus the logarithm of the DNA concentration (orange dots). The data was taken from Garbett, Nichola C. *et al.*, 2007. The fitted curve assuming a one-to-one binding model is shown as a black line.

**Two-state unfolding of the Arc dimer**

The unfolding reaction of the Arc repressor was explained by a two-state transition from folded dimer to unfolded monomer ($N_2 \rightleftharpoons 2U$) by Bowie and Sauer, 1989 (4). We globally fitted the normalised fluorescence data (digitised from Figure 3a from ref. (4)) yielding parameters similar to those previously reported (Figure S2). We obtained an *m*-value of 2 kcal/mol/M, compared to the published *m*-value of 1.9 kcal/mol/M.

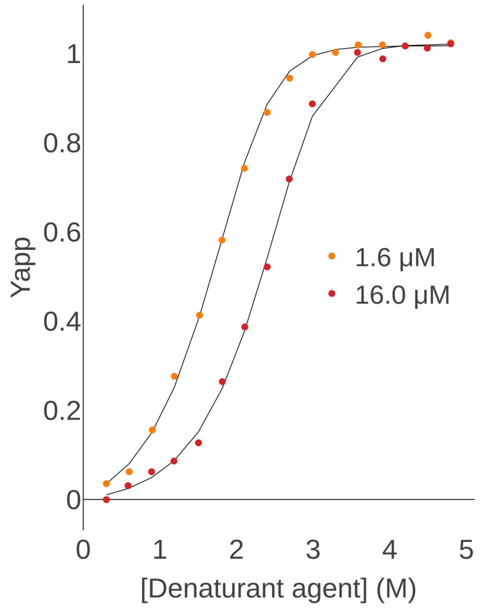


**Figure S3.** Apparent unfolded fraction of the Arc repressor as a function of urea concentration at two different protein concentrations. Adapted from (4). B) ChiraKit based two-state ($N_2 \rightleftharpoons 2U$) global fitting of the data extracted from the Figure in panel A.

#### Three-state unfolding of the FtsZ dimer

FtsZ, a major protein in bacterial cytokinesis that polymerizes into single filaments, unfolds in urea via a dimeric intermediate ($N_2 \rightleftharpoons I_2 \rightleftharpoons 2U$), as presented by Montecinos-Franjola *et al.*, 2012 (5). The normalised CD data presented in Figure 3A was digitised and globally fitted, obtaining comparable results (Figure S3): 6.4 versus 5.9 kcal/mol/M for the $m_1$-value, and 3.2 versus 2.4 kcal/mol/M for the $m_2$-value.

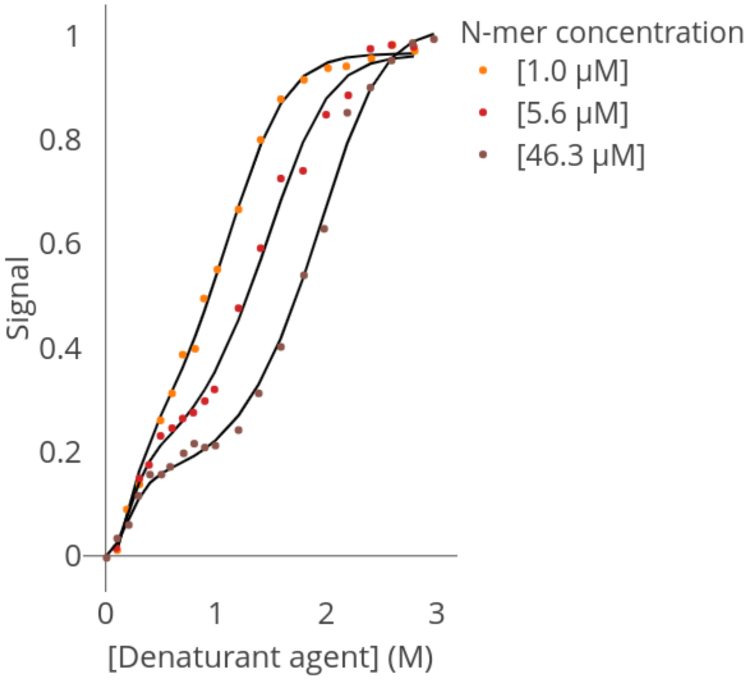


**Figure S4.** Apparent unfolded fraction of FtsZ as a function of urea concentration at three different protein concentrations (circles: 1 μM, triangles: 5.6 μM, and squares: 46.3 μM). Adapted from (5). B) ChiraKit based three-state ($N_2 \rightleftharpoons I_2 \rightleftharpoons 2U$) global fitting of the data extracted from the Figure in panel A.

#### Supplementary Methods

#### Fitting of the Protein - DNA binding curves

The string used to estimate the equilibrium dissociation constant ($K_D$) of the interaction between berenil and calf thymus DNA in the 'Custom analysis' panel was:

'0.5*(ABsignal - freeAsignal) * ((10**logTenOfKd+6+Bconc) - sqrt((10**logTenOfKd+6+Bconc)**2 - 4*6*Bconc)) + freeAsignal*6'

It represents the Equation:

$$Y = 0.5(C - A) * (10^x + 6 + [DNA] - \sqrt{(10^x + 6 + [DNA])^2} + 6A \quad (3)$$

where $C$ and $A$ weight, respectively, the complex and free DNA contribution to the signal. The parameter to be fitted is $x$ ($10^x$ equals the $K_D$).
